# Density dependence in demography and dispersal generates fluctuating invasion speeds

**DOI:** 10.1101/075002

**Authors:** Lauren L. Sullivan, Bingtuan Li, Tom E. X. Miller, Michael G. Neubert, Allison K. Shaw

**Author notes:** Author Contributions: T.E.X.M., M.G.N., and A.K.S. designed initial research; L.L.S. and M.G.N. led simulations; L.L.S. led writing; All authors contributed to ideas development and writing.

## Abstract

Density dependence plays an important role in population regulation and is known to generate temporal fluctuations in population density. However, the ways in which density dependence affects spatial population processes, such as species invasions, are less understood. While classical ecological theory suggests that invasions should advance at a constant speed, empirical work is illuminating the highly variable nature of biological invasions, which often exhibit non-constant spreading speeds even in simple, controlled settings. Here, we explore endogenous density dependence as a mechanism for inducing variability in biological invasions with a set of population models that incorporate density dependence in demographic and dispersal parameters. We show that density dependence in demography at low population densities—i.e., an Allee effect—combined with spatiotemporal variability in population density behind the invasion front can produce fluctuations in spreading speed. The density fluctuations behind the front can arise from either overcompensatory population growth or from density-dependent dispersal, both of which are common in nature. Our results demonstrate that simple rules can generate complex spread dynamics, and highlight a novel source of variability in biological invasions that may aid in ecological forecasting.

## Introduction

Fluctuations in population size have long fascinated ecologists and fueled a now-classic debate over whether populations are governed by extrinsic environmental factors or by intrinsic self-limitation (15). One of the most important advances of twentieth-century ecology was the discovery that intrinsic density feedbacks can cause population densities to fluctuate, even in constant environments (26; 5; 48). This discovery helped resolve the important role of density dependence in population regulation, revealing that strong regulating forces can generate dynamics superficially consistent with no regulation at all. Our understanding of temporal fluctuations in population size stands in sharp contrast with our relatively poor understanding of fluctuations in the spatial dimension of population growth: spread across landscapes.

Understanding the dynamics of population spread takes on urgency in the current era of human-mediated biological invasions and range shifts in response to climate change. The velocity of spread, or “invasion speed”, is a key summary statistic of an expanding population and an important tool for ecological forecasting (8). Estimates of invasion speed are often derived from regression methods that describe change in spatial extent with respect to time (30; 1; 49). Implicit in this approach is the assumption that the true spreading speed is constant and deviations from it represent “error” in the underlying process, or in human observation of the process. This assumption is reinforced by long-standing theoretical predictions that, under a wide range of conditions, a population will asymptotically spread with a constant velocity. Invasion at a constant speed can arise from both *pulled* waves (where the advancing wave moves forward by dispersal and rapid growth of low-density populations far in front of the advancing wave (56; 44; 16; 32)), as well as *pushed* waves (where the invasion is driven by reproduction and dispersal from high-density populations behind the invasion front (21; 55; 50)). The conventional wisdom of a long-term constant invasion speed is widely applied (53; 9).

In contrast to classic approaches that emphasize a long-term constant speed, there is growing empirical recognition that invasion dynamics can be highly variable and idiosyncratic (27; 29; 34; 59; 60; 4; 54; 14). There are several theoretical explanations for fluctuations in invasion speed (which we define here as any persistent temporal variability in spreading speed), including stochasticity in either demography or dispersal (35; 54; 17; 42; 14), and temporal or spatial environmental heterogeneity (43; 33; 57; 58; 3; 40). Indeed, empirical studies often attribute temporal variation in speed to differences in the environments encountered by the invading population (e.g., (1; 37)). Predator-prey dynamics can also induce fluctuating invasion speeds (33; 7). Notably, Dwyer and Morris (7) showed that density feedbacks can produce fluctuations in spreading speed, yet we still have an incomplete understanding of the conditions under which fluctuations in speed arise. Surprisingly few theoretical studies have since investigated these density feedbacks, especially with respect to their effect on endogenously-driven speed fluctuations, despite recent empirical work on invasion variability (34; 59; 53; 60).

Here, we develop deterministic, single-species mathematical models of spatial spread to ask under what conditions the invasion speed of an expanding population can fluctuate in a spatially uniform and temporally constant environment. As a starting point, we took inspiration from the relatively complete understanding of fluctuations in population size generated by density dependence in nonspatial models (48). We conjectured that density-dependent feedbacks might similarly generate fluctuating invasion speeds pursuing the suggestion first made in (7). Because spread dynamics are jointly governed by demography (local births and deaths) and dispersal (spatial redistribution), we considered several types of density feedbacks (39), including positive density dependence in population growth (i.e., Allee effects) at the low-density invasion front (47), and density-dependent movement (25; 7).

Our analysis uncovered novel density-dependent mechanisms that can induce variability in invasion speed, with fluctuations ranging from stable two-point cycles to more complicated aperiodic dynamics. By demonstrating that simple invasion models can generate complex spread dynamics, our results reveal previously undescribed sources of variability in biological invasions and provide a roadmap for empirical studies to detect these processes in nature.

## Models and Results

We use integrodifference equations (16) to model population growth and spread. These models describe the change in population density (*n*_*t*_(*x*)) from time *t* to time *t* + 1 as the result of demography and dispersal. First, individuals at location *y* generate *f*(*n*_*t*_(*y*)) offspring and then die. Next, a fraction *p* of these offspring disperse. The probability that a dispersing individual moves from location *y* to location *x* is given by the dispersal kernel, *k*(*x – y*). The remaining fraction (1 – *p*) remain at their natal location. Concatenating reproduction and dispersal, we have (51; 52; 22; 23):

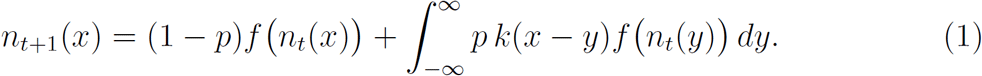
We will assume that *f*(1) = 1, so that the population has an equilibrium at the carrying capacity *n*_*t*_(*x*) = 1, and that the tails of the dispersal kernel *k* are thin (i.e., go to zero at least exponentially fast), so that the probability that an individual disperses an extremely large distance is exceedingly small.

In general, both the dispersing fraction *p* and the dispersal kernel *k* may depend on the population density at the natal location, as does the reproduction function *f*. The way that the functions *f, p*, and *k* depend on population density determine the dynamics of Eq. 1. In the simplest case, the reproduction function *f* is strictly compensatory; that is, *f* is an increasing but decelerating function of density (*f*′(*n*) > 0 and *f*″(*n*) < 0). For strictly compensatory models, the population will spread at a constant asymptotic speed (Fig. 1a) if three conditions hold: small populations grow (*f*′(0) > 1), all individuals disperse (*p* = 1), and dispersal distance is independent of population density. Here, the speed is determined by the growth and spread of the low-density populations far ahead of the main invasion front (56); the dynamics at high densities do not matter – the hallmark of a pulled invasion.

**Figure 1:**
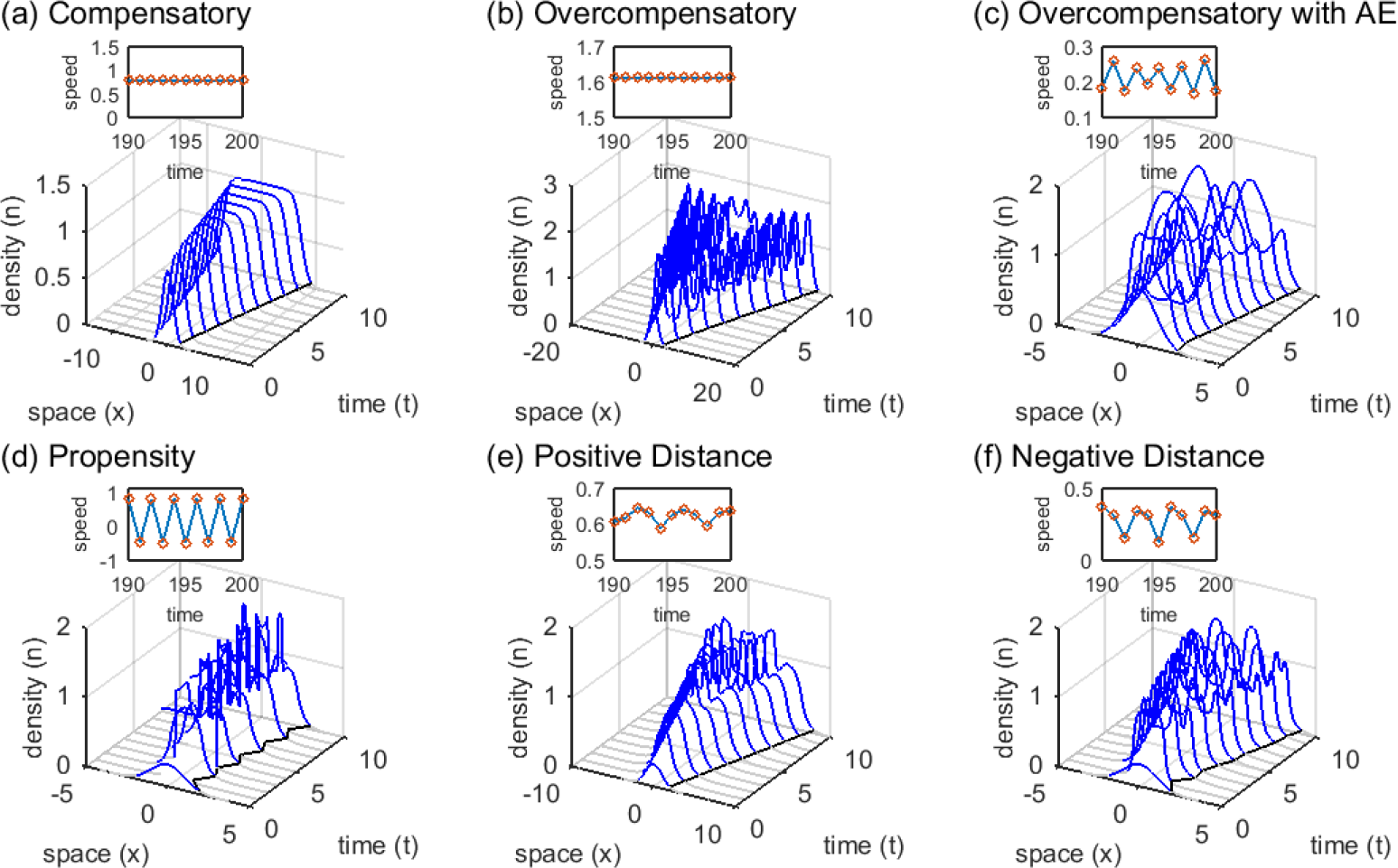
Invasion dynamics under different types of density dependence and dispersal. With compensatory growth at high densities (a), the wave shape and invasion speed are both constant. This is true with and without low-density Allee effects (overcompensatory model: *σ*^2^ = 0.25, *a* = 0, and *r* = 0.9; Fig. S1a). With over-compensatory population growth and no Allee effect (b), population density exhibits fluctuations behind the front yet the leading edge progresses at a constant speed (overcompensatory model: *σ*^2^ = 0.25, *a* = 0, and *r* = 2.7; Fig. S1a). However, when overcompenstion combines with low-density Allee effects (c), the invasion speed fluctuates (overcompensatory model: *σ*^2^ = 0.25, *a* = 0.4, and *r* = 2.7; Fig. S1a). Variability in invasion speed can also occur when Allee effects combine with density-dependence in the proportion of dispersing offspring (d) (propensity model: *a* = 0.2, *λ* = 0, 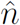 = 0.9, *p*_0_ = 0.05, *p*_*max*_ = 1, *α* = 50), or in dispersal distance (e,f). In the latter model, dispersal distance decreases with population density (e) (distance model: *a* = 0.2, *λ* = 0, 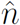 = 0.9, *β* = −50, 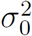, 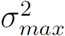), or increases with density (f) (distance model: parameters as in (e) except *β* = 50). Initial population densities are either 2 (a-c) or 0.8 (d-f) times the standard normal probability density truncated at |*x*| = 5.

Constant asymptotic invasion speeds are not, however, limited to the simple case just described. In the absence of Allee effects, they can also occur if the reproduction function produces overcompensation—declining offspring production with increasing population density (so that *f*′(1) < 0). As with classic non-spatial models, over-compensation produces oscillations in population density (26; 5; 48), which in turn cause dynamic changes in the shape of the wave behind the invasion front. Despite these complex fluctuations at high population densities, the invasion speeds of over-compensatory models (without Allee effects) remain constant (Fig. 1b), and are still determined by the dynamics at low densities (19).

Long-standing theory suggests that invaders subject to Allee effects at low population density and compensatory dynamics at larger population density, will also eventually spread at a constant speed if their initial population sizes are sufficiently large and the Allee effect is not too strong (55; 21). Allee effects cause invasion waves to be pushed from behind their leading edge (16; 55). When Allee effects are sufficiently strong, the invasion speed no longer depends upon the pull of populations at low densities in front of the wave, but rather on the strength of the push from the high density populations behind it. In our models, we show that when low-density Allee effects combine with spatiotemporal population density fluctuations (created through overcompensation or density-dependent dispersal), the invasion speed may not be constant asymptotically, as expected under classic invasion theory, but may rather exhibit persistent fluctuations (Fig. 1c-f).

### Allee effects and overcompensation

First, we investigated whether combining an Allee effect with overcompensation at high population density could induce fluctuating invasion speeds when dispersal is density-independent and all offspring disperse (i.e., *p* = 1). This model (the ‘over-compensatory model’, see Materials and Methods, Fig. S1a) has two important parameters: *r*, which affects both the growth rate at low density and the strength of density dependence at carrying capacity, and *a*, the Allee threshold. We assume that when the population density falls below *a*, no offspring are produced there (a strong Allee effect). If the population density falls below *a* everywhere, the population is doomed to extinction.

Simulations (described in Materials and Methods) revealed this model generates variable-speed invasions (Fig. 1c), but only when the low-density Allee threshold is of intermediate value and high-density overcompensation is strong (*r* > 2, Fig. 2a). For *r* > 2, the local equilibrium density *n*_*t*_(*x*) = 1 is unstable, leading to sustained fluctuations in local density. Our simulations suggest *r* > 2 is a necessary condition for fluctuating invasion speeds in the overcompensatory model. If the Allee threshold (*a*) is too large, the spreading population eventually falls below the threshold everywhere and is extirpated. If *a* is sufficiently small, the invasion proceeds with an apparently constant speed (Fig. 2a).

**Figure 2:**
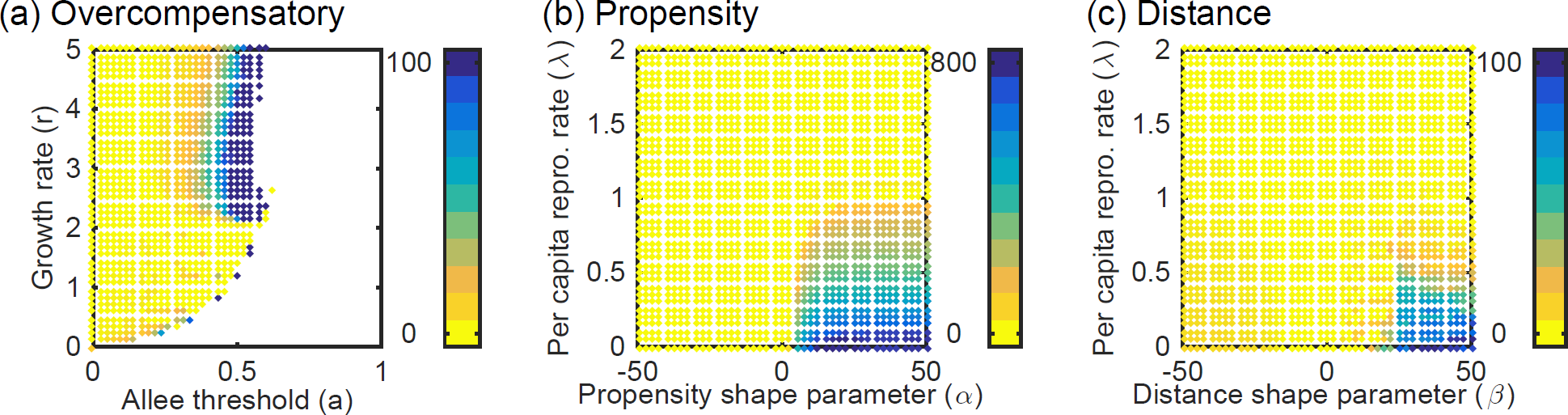
Amplitude of fluctuations in invasion speed normalized by the mean speed for populations with Allee effects and (a) overcompensatory growth, (b) density-dependent dispersal propensity, and (c) density-dependent dispersal distance. Darker colors (blue) indicate where fluctuations create large differences from the mean invasion speed. Values of zero (yellow) indicate invasion waves that move at a constant speed. White regions indicate where invasions fail. Parameter values: *σ*^2^ = 0.25 (a-b); *a* = 0.2, 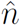 = 0.9 (b-c); *p*_0_ = 0.05, *p*_*max*_ = 1 (b); 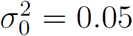, 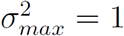; Initial population densities are equivalent to those in Fig. 1.

These fluctuations are induced by the combination of a strong Allee effect, which produces a pushed wave, and strong overcompensation, which produces large spatiotemporal variation in density behind the invasion front and thus variation in the strength of the push (Fig. 3). When the population density at any location is smaller than the Allee threshold (*a*), as at the leading edge of the wave, the population vanishes before the next time step. Populations just above *a* become large after reproduction, but as the population size increases beyond *a*, the offspring population size *f*(*n*(*x*)) declines as a result of overcompensation (Fig. S1a). Therefore, when reproduction occurs (transition between *n*(*x*) and *f*(*n*(*x*)), Fig. 3 black vs blue), populations with the highest density become populations of low density, and populations with density just above *a* become high density. Through time, this creates variability in the size of the push by varying the size of the region contributing to the wave front, leading to fluctuating invasion speeds (Fig. 3d, S3a-f). The speed fluctuations can be periodic or more complex (Fig. S2). They vary in amplitude by as much as 100% of the mean speed, with some parameter combinations reaching amplitudes of *∼* 400% of the mean speed (Fig. 2a).

This mechanism for variable-speed invasion does not depend on the discreteness of time. We developed a continuous-time version of the overcompensatory model, where we find fluctuating invasion speeds as long as density fluctuations behind the wave front combine with strong low-density Allee effects (SI Appendix, Fig. S4-6).

**Figure 3:**
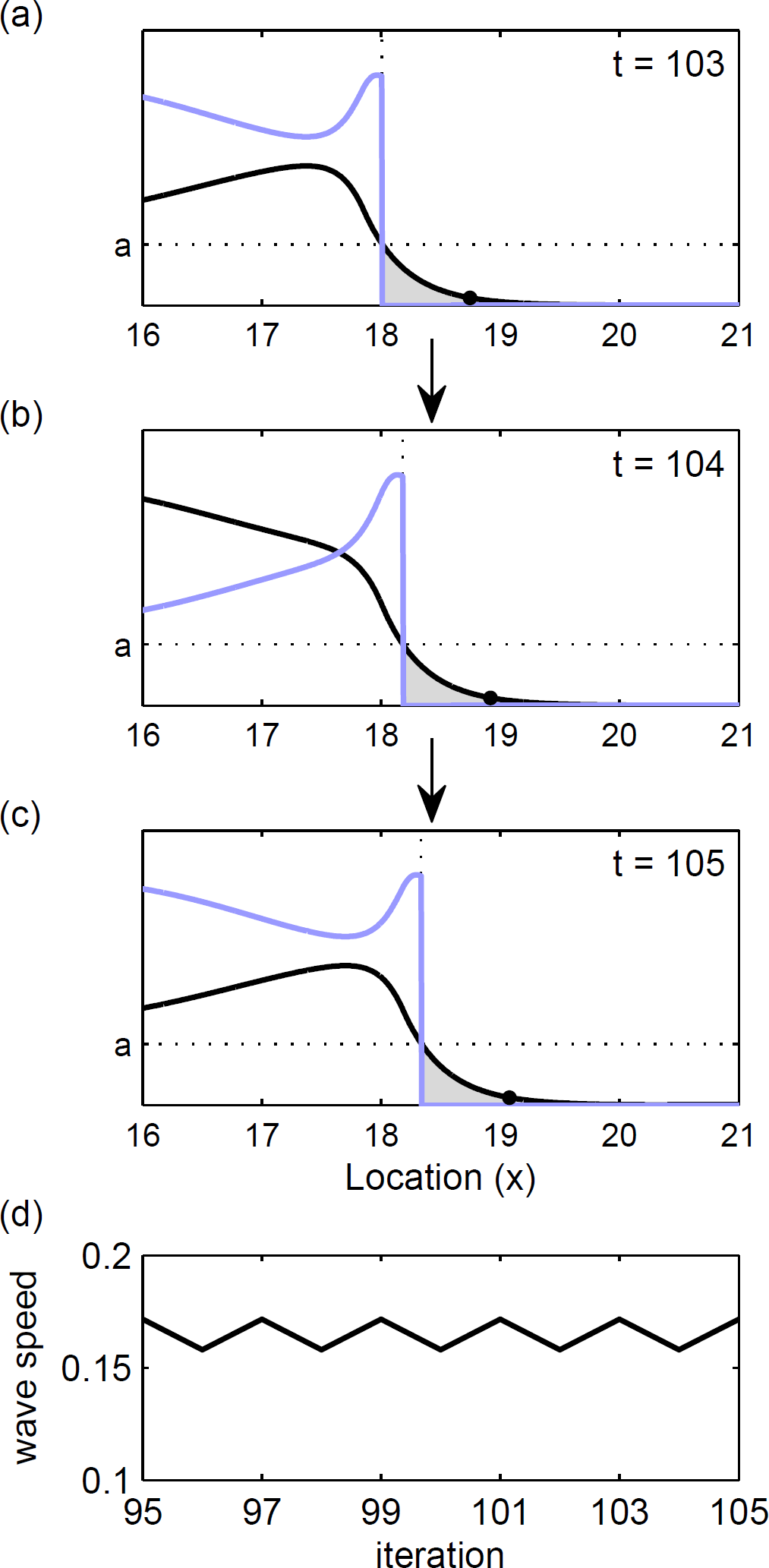
Population density before (*n*(*x*) - black curve) and after (*f*(*n*(*x*)) - blue curve) growth (overcompensatory model with Allee effects, Eq. 3) at sequential time steps (a-c). Gray regions represent locations that go extinct due to Allee effects (light gray; *n*(*x*) < *a*), and the solid point shows the edge of the wave. The wave speed over time (d), corresponds to (a-c). Parameter values include *r* = 2.2, *a* = 0.4, *σ*^2^ = 0.25.

### Allee effects and density-dependent dispersal

Overcompensation is not the only mechanism that can generate the spatiotemporal variability in population density that is necessary to produce fluctuating invasion speeds when combined with Allee effects. Density-dependent dispersal, manifest as either density-dependence in the propensity to disperse (*p*) or in the shape of the dispersal-kernel (*k*), can generate this high-density variability in the pushing force as well. We demonstrate this result with two models (the ‘propensity model’ and the ‘distance model’, respectively, see Materials and Methods), both built upon a piecewise linear growth function that is compensatory at high population density (Fig. S1b). We continue to include low-density Allee effects. When the population size falls below the threshold density *a*, individuals produce offspring at the constant per capita rate *λ*. Alternatively, if the population size exceeds *a*, the population goes to carrying capacity.

In the propensity model, population density influences the propensity to disperse (*p*). In particular, we assume that the proportion of offspring that disperse is given by a logistic function of local population density (*n*_*t*_(*x*)) (Eq. 5) with four parameters: the minimum (*p*_0_) and maximum (*p*_max_) dispersal proportion; a location parameter 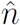, which is the density at which the dispersal propensity is halfway between *p*_0_ and *p*_max_; and a shape parameter *α*. The sign of *α* determines if the proportion dispersing increases (*α* > 0) or decreases (*α* < 0) with density (Fig. S1c). The larger the magnitude of *α* the steeper the density response, which is centered around 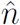.

The propensity model can also generate invasions that spread at fluctuating speeds (Fig. 1d, S7). We found these fluctuations persist only when Allee effects are strong (0 ≤ *λ* < 1), dispersal propensity increases with population density (*α* > 0), and the dispersal response occurs at a population density that is larger than the Allee threshold (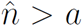. Fluctuations in speed are nearly always periodic (Fig. S7c, S8a-d) and of large amplitude, altering the invasion speed by ∼ 100% − 750% relative to the mean speed (Fig. 2b). These large-amplitude periodic fluctuations often include positive and negative speeds, meaning that invasions alternate between steps forward and smaller steps backward (Fig. 1d).

As before, spreading speed fluctuations are created through variations in the dispersing population that pushes the invasion forward from behind the front (Fig. 1d). The magnitude of the push depends on the width of the region contributing dispersing individuals, and the proximity of this region to the front (Fig. S3g-l). When density dependence in dispersal is strong and positive (large *α*), the population directly adjacent to the front is below the Allee effect threshold (*a*) and therefore decays to zero (Fig. S3g-h). Farther behind the front, density is above *a*, but below the dispersal midpoint (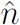), thus this region of the population reproduces but does not disperse (Fig. S3h-i). This action results in a large push from behind the wave front that moves the invasion forward at the next time step when the non-dispersing population eventually disperses (Fig. S3i-k). Subsequently, the region of the non-dispersing population is much smaller and farther from the invasion front at the next time step, resulting in a much smaller push (Fig. S3k).

With the distance model we explore a second type of density-dependent dispersal, where density alters the dispersal distance. Here, all offspring disperse (*p* = 1), but density alters the variance (*σ*^2^) of the dispersal kernel (Eq. 6). Four parameters control this dependence: 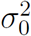 and 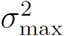, which are the lower and upper bounds of the variance; the location parameter, 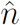 which is the density at which dispersal variance is halfway between 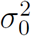 and 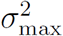; and a shape parameter *β*. The dispersal variance increases with population density when *β* is positive, and decreases with density when *β* is negative. The larger the absolute value of *β*, the sharper the response (Fig S1d).

The distance model also produces the necessary spatiotemporal variability in population density behind the invasion front to induce fluctuating invasion speeds (Fig. 1e,f, S7). As in the propensity model, the invasion speed only fluctuates when Allee effects are strong (0 ≤ *λ* ≤ 1). However, unlike the propensity model, we find persistent fluctuations are possible when density-dependent dispersal is both positive (*β* > 0) and negative (*β* < 0) (Fig. 2c). The speed fluctuations are more frequently aperiodic (Fig. S8e-h) than the two-cycle fluctuations seen in the propensity model, with largest amplitude when dispersal distance increases with density (*β* > 0) (Fig. 2c, S7f). In general, fluctuations are larger as both Allee effects and density-dependent dispersal are stronger, and alter the invasion speed by *∼* 5% *−* 100% (*β* > 0), and *∼* 1% − 9% (*β* < 0) relative to the mean speed (Fig. 2c, S7f).

When the dispersal distance exhibits strong positive density dependence (Fig. S3m-r), populations at densities above the dispersal threshold disperse long distances, and those below disperse short distances. In this model, each push forward is made up of a combination of both short and long distance dispersers. The size of this push changes depending on the proportion of the push made up of each type of disperser, which is temporally variable, creating fluctuating invasion speeds. A similar mechanism operates when *β* < 0 (Fig. S3s-x), however instead high density populations disperse short distances and vice versa.

## Discussion

Our work provides novel insight into mechanisms behind invasion variability: fluctuations in invasion speed can occur solely due to endogenous density dependence. In the models we examine, both a strong low-density Allee effect (creating a pushed wave (9; 28)), and large variations in population density behind the invasion front are necessary to create fluctuating invasion speeds. We demonstrate that the necessary spatiotemporal variability can be generated via two types of density feedbacks: overcompensatory density dependence, or density-dependent dispersal. When combined with Allee effects, either of these factors can cause the strength of the invasion push from high density populations to vary, leading to varying spreading speeds. The potential for deterministic, density-dependent processes to generate complex fluctuations in local population density is a canonical result of theoretical population biology (15; 26; 5; 48) and has proven influential in basic and applied empirical settings (36). By considering the spatial dimension of population growth, which is increasingly relevant in the context of global change, our new results flesh out understanding of complex population dynamics arising from endogenous mechanisms. We conjecture that there is some generality to this mechanism as we also see fluctuating speeds in continuous time (SI Appendix, Fig. S4), although we recognize fluctuations can occur through other means (e.g. (7; 33; 14)). Our results are potentially consistent with the highly variable spreading speeds seen in empirical invasion studies (14; 34; 59; 54; 4; 60).

Processes capable of generating fluctuations in population density that create the variable pushing force behind the invasion vanguard are common in nature. First, many invasive species show the combination of high intrinsic growth rates and con-specific interference at high density that gives rise to overcompensatory population fluctuations (36; 61). Second, density dependent dispersal as a distinct source of spatiotemporal density fluctuations ions can arise even with strictly compensatory density dependence in population growth. We found fluctuating invasion speeds with positive density-dependent dispersal propensity, which is common in organisms with environmentally inducible dispersal polymorphisms, including many insects. For example, wingless aphids (11; 13) and planthoppers (38) can produce winged morphs when densities become high. When density dependence alters dispersal distance, fluctuations in speed were seen under both positive and negative density dependence. Mobile organisms can increase their dispersal distance with increasing density by altering behavioral responses (25). Alternatively, dispersal distances can decrease with density when crowding decreases reproductive and dispersal ability (24; 6; 25), or in animals (notably small mammals) with strong group behavior (12; 2; 25).

Allee effects, a common density-dependent process (18; 31), influence small populations by decreasing low-density vital rates (e.g., reproduction (51)). We find in all of our models that Allee effects, and the pushed invasions that they generate, are a necessary ingredient of fluctuating speeds. Interestingly, this result contrasts with Dwyer and Morris (7). Working with a two-species model, they found that fluctuating speeds can occur when predator dispersal distance depends on prey density (a type of density dependent movement) but without an explicit Allee effect. We conjecture that predator-prey dynamics in their model may in fact give rise to an implicit Allee effect, as is known to occur in other predator-prey models (33). Biologically, density-dependent movement can contribute to an Allee effect by reducing mate finding abilities at low densities, especially when the movement is sex biased (53; 41). In this way, the study by Dwyer and Morris (7), while superficially inconsistent with ours, may nonetheless satisfy the conditions we identify as necessary for variable invasions.

Thoroughly accounting for the sources of variability in the speed of biological invasions may improve invasion forcasting. Our work suggests that intrinsic density dependence can create complex invasion dynamics, consistent with the highly variable spreading speeds seen in empirical invasion studies (34; 59; 60; 14; 4; 54). However, it remains an open question whether and how often these processes affect the ecological dynamics of spread, given the pervasive influences of environmental heterogeneity (43; 33; 57; 58; 3; 40) and demographic stochasticity (35; 17; 42), and their roles in invasion variability. To begin to answer this question, we suggest coupling models and empirical data, which has proven to be a fruitful approach to understanding the intrinsic mechanisms behind fluctuations in local population density (e.g., (5; 48)). Collecting long-term data can be difficult, but some patterns might be straightforward to identify from existing datasets. In particular, the strong two-cycle speed fluctuations generated when invaders experience both Allee effects and density-dependent dispersal propensity would likely be detectable in data. Few empirical studies have tested for endogenous mechanisms of fluctuating invasion speeds, including studies for which variability in speed was an explicit focus (53; 27; 29; 34; 59) (but see (14)). Thus, signatures of endogenous variability may be embedded in existing data, and we encourage empiricists to re-examine variable invasion data in the context of these density-dependent mechanisms.

## Materials and Methods

The models we studied are each a special case of equation (1). They all use the Laplace dispersal kernel with variance *σ*^2^:

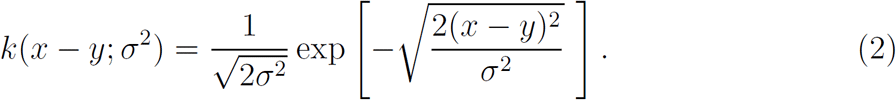

Qualitative results are robust to kernel choice (i.e. Normal, Cauchy).

### Overcompensatory Model

We combine low-density Allee effects with the possibility of overcompensation at high density (Fig. S1a):

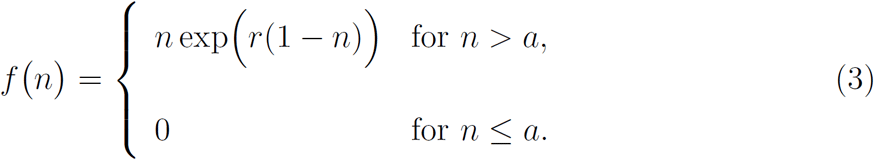

Dispersal is independent of density in this model (*σ*^2^(*n*) = *σ*^2^, a constant) and all offspring disperse (*p* = 1).

### Propensity Model

Here, we used a linear-constant model for growth 
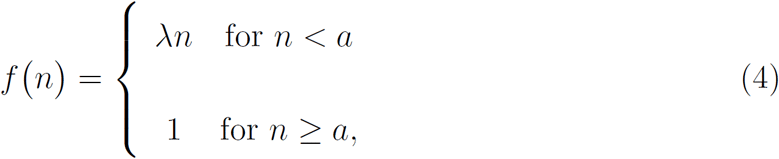
where 0 ≤ *a* < 1 (Fig. S1b). Dispersal propensity depends upon the population density (*n*_*t*_(*x*)) via a logistic form similar to other models with density-dependent dispersal (Fig. S1c) (45):

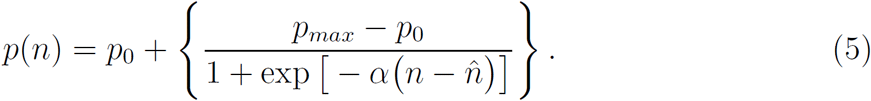

As in the overcompensatory model, the distance moved by dispersing individuals is independent of density (*σ*^2^(*n*) = *σ*^2^, a constant).

### Distance Model

For this model, we use the reproduction function (4), but assume all offspring disperse (*p* = 1) following a dispersal distribution whose variance is a logistic function of parental density (*n*_*t*_(*x*)) (Fig. S1d). I.e., 
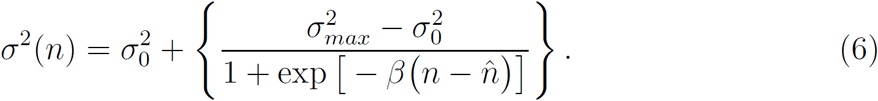

We simulated each model for 200 iterations across a domain of length 1200 with 2^16^ + 1 spatial nodes. Within each simulation, we defined the location of the invasion front at each time step as the location where the density of the invasion wave first exceeded a density threshold of 0.05. We then used this location to calculate: (1) the instantaneous invasion speed (i.e., the distance traveled by the front between consecutive time steps), (2) the mean invasion speed averaged over the last 50 time steps, and (3) the amplitude of invasion speed fluctuations (the difference between the maximum and minimum speed over the last 20 time steps). See Table S1 for a list of parameters and definitions. Code to run these models and recreate all figure will be available at Dryad upon manuscript acceptance.

**Table 1:** All model parameters, definitions and corresponding models.

| Variable | Meaning |
| --- | --- |
| $t$ | time |
| $x, y$ | locations |
| $n_t(x)$ | population density at location $x$ and time $t$ |
| Parameter | Meaning |
| $a$ | Allee effect threshold |
| $r$ | intrinsic growth rate (overcompensatory model) |
| $\lambda$ | low-density per capita reproductive rate (propensity and distance models) |
| $\hat{n}$ | dispersal density midpoint parameter (propensity and distance models) |
| $p$ | fraction of offspring that disperse |
| $p_0$ | minimum dispersal propensity (propensity model) |
| $p_{max}$ | maximum dispersal propensity (propensity model) |
| $\alpha$ | propensity shape parameter (propensity model) |
| $\sigma^2$ | variance of the dispersal kernel |
| $\sigma_0^2$ | minimum dispersal variance (distance model) |
| $\sigma_{max}^2$ | maximum dispersal variance (distance model) |
| $\beta$ | distance shape parameter (distance model) |
| Function | Meaning |
| $k(x - y)$ | dispersal kernel |
| $f(n_t(x))$ | growth or offspring density |

## Acknowledgements

LLS and AKS were supported by startup funds from the University of Minnesota (UMN) to AKS, BL by NSF DMS-1515875, TEXM by NSF DEB-1501814, and MGN by NSF DEB-1257545 and DEB-1145017. The initial idea was developed during the 2014 ACKME Nantucket Mathematical Ecology retreat with input from participants and funding from WHOI Sea Grant. The manuscript was greatly improved by comments and support from E. Strombom, R. Williams, an anonymous editor, and two anonymous reviewers. The UMN Minnesota Supercomputing Institute (MSI) provided resources that contributed to the research results reported within this paper. URL: http://www.msi.umn.edu.

## Supporting Information (SI)

**Figure S1:**
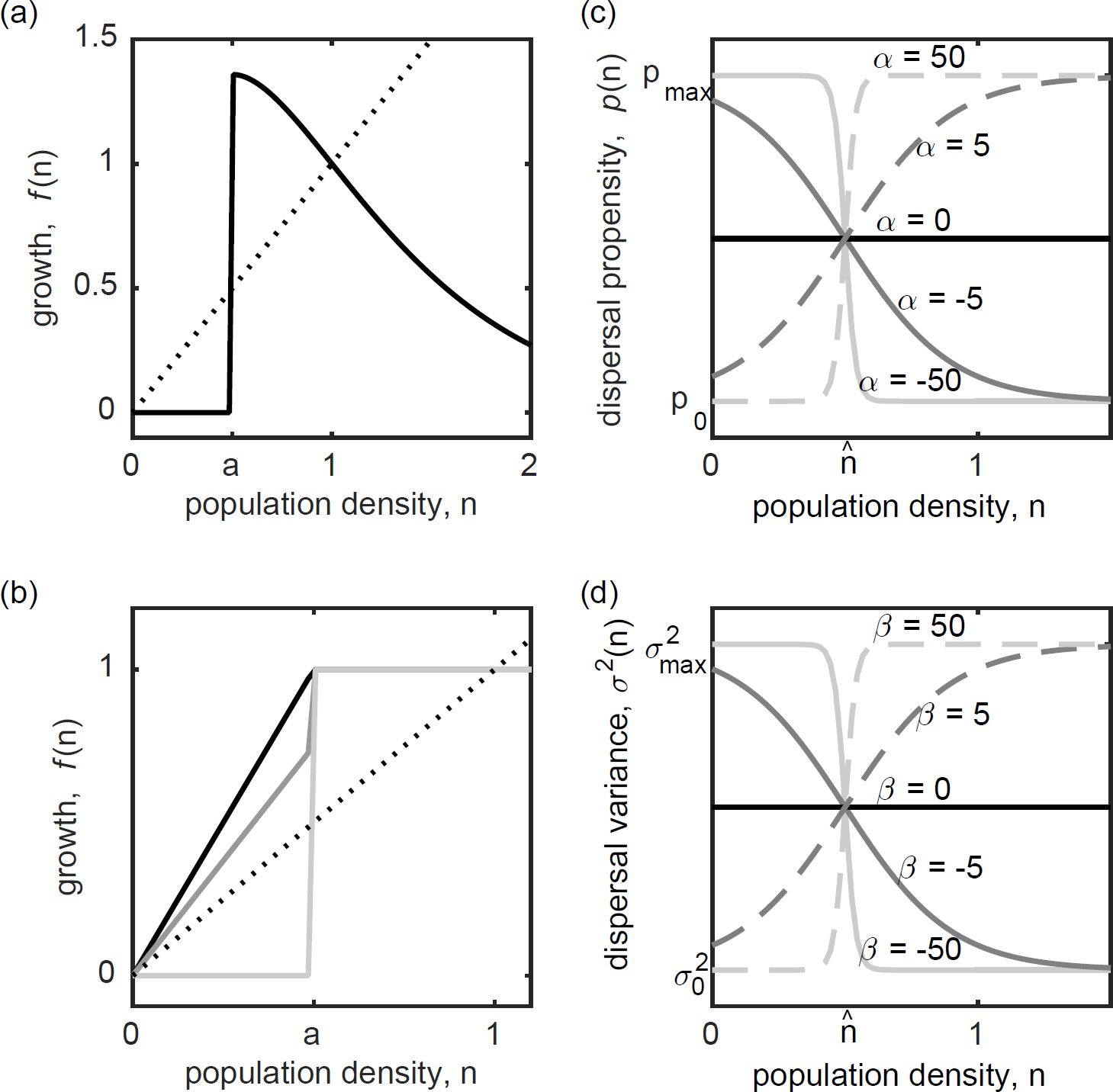
Reproduction and dispersal functions used in the overcompensatory, propensity, and distance models (described in Materials and Methods). (a) The reproductive rate f(n) as given by Eq. 3 where *r* = 0.9 and *a* = 0 (same parameterization as Eq. 3 for Fig 1a, black), *r* = 2.7 and *a* = 0 (same parameterization as Eq. 3 for Fig 1b, dark gray solid), *r* = 2.7 and *a* = 0.4 (same parameterization as Eq. 3 for Fig 1c, light gray dashed). (b) The reproductive rate f(n) as given by Eq. 4 when *a* = 0.5 and *λ* = 0 (light gray), *λ* = 1.5 (dark gray), *λ* = 2 (black). Parameterization here is close to that of Fig 1d-f, except *a* is smaller here to more clearly visually demonstrate the differences between strong, weak and no Allee effects. (c) The propensity to disperse when altered by density dependence as given by Eq. 5 for different *α*. (d) The variance of the dispersal kernel when altered by density dependence Eq. 6 for different *β* values.

**Figure S2:**
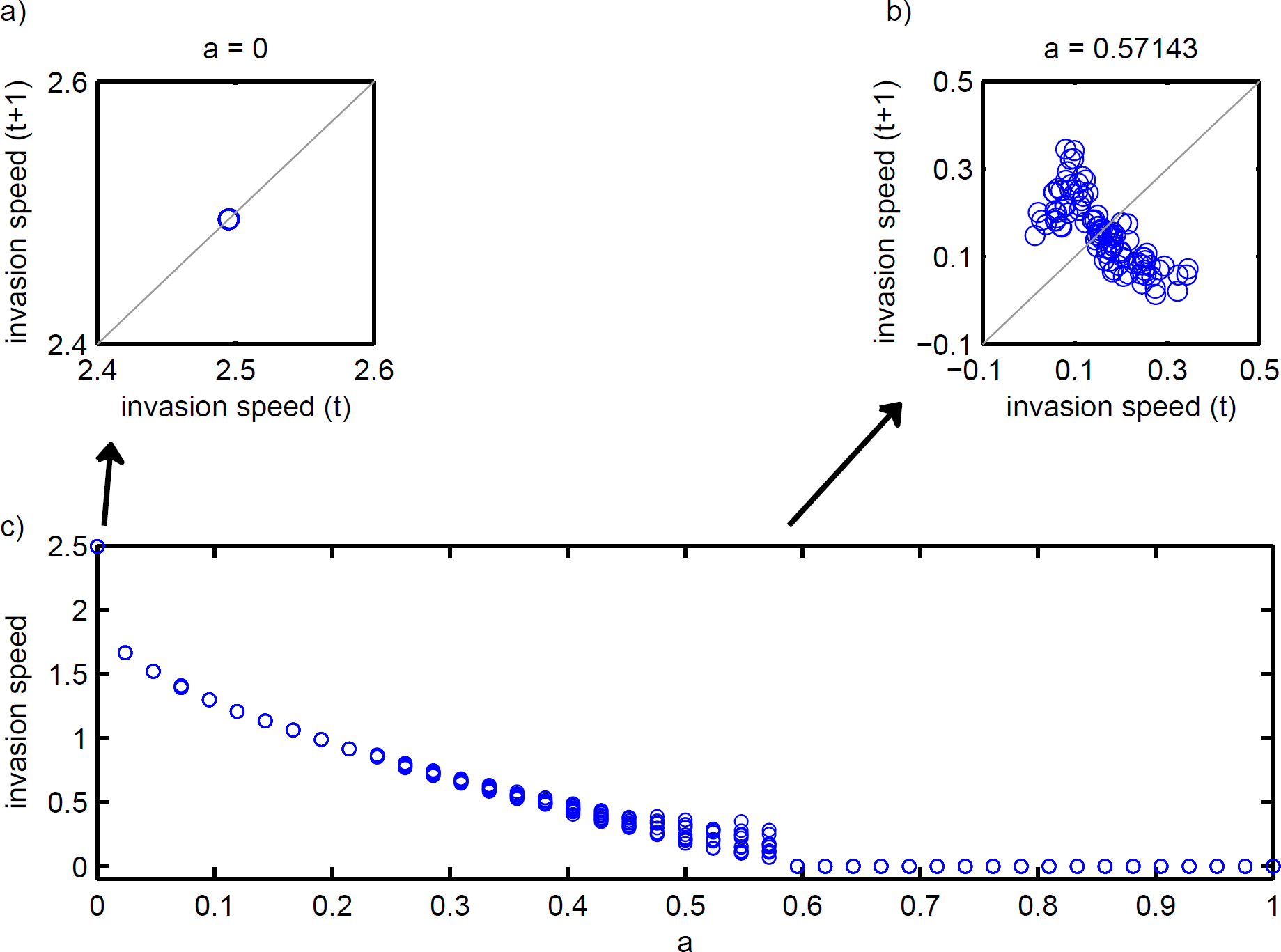
The periodicity of the invasion speed through time for the overcompensatory model - Allee effects and overcompensation. In panels a-b, the wave position is plotted at time *t* vs time *t* + 1. The wave speed ranges in periodicity across values of the Allee effect threshold *a*. At small values of *a* the invasion speed is constant (a), and at larger *a* values (b), the wave speed becomes chaotic until *a* becomes so large the population goes extinct. In panel (c), the range of invasion speeds represents the amplitude of fluctuations. At each plotted *a* value, the invasion speed for the previous 100 time steps are plotted. When points appear as hollow points, the same invasion speed is being plotted over itself many times. Here, *σ*^2^ = 0.25.

**Figure S3:**
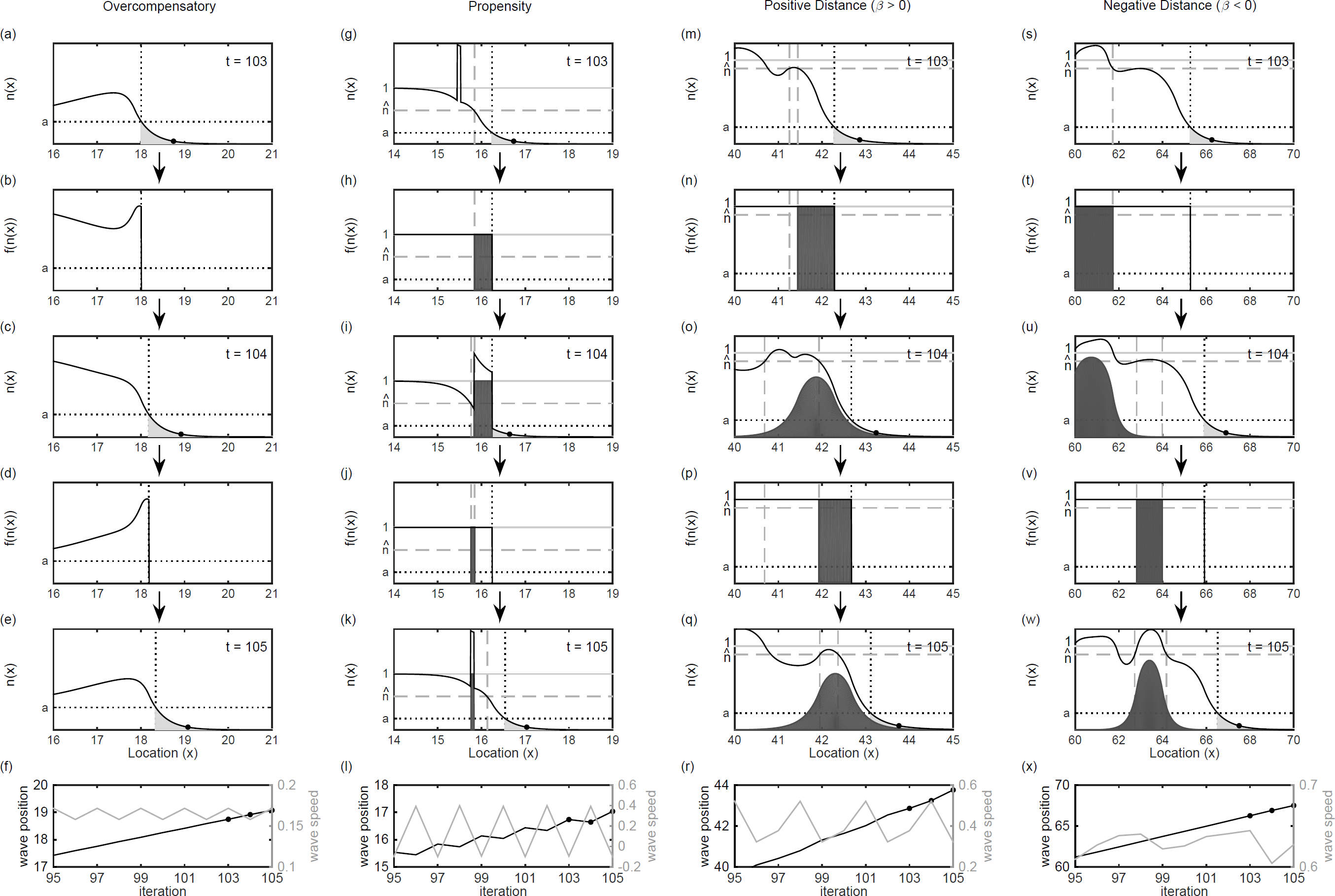
Four examples of fluctuations in invasion speed. The top five rows show the population density before (*n*(*x*)) and after (*f*(*n*(*x*))) growth at sequential time steps, showing individuals that will not reproduce (light gray; *n < a*), those that do not disperse far (dark gray; *n* > 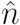 or *n* < 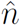), and the edge of the wave (solid point). The bottom row shows the wave position and speed over time. Parameter values and initial densities are the same as Fig. 2 except: (a-f) *r* = 2.2, *a* = 0.4, (g-l) 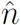 = 0.6, *α* → ∞, (m-r) *β* → ∞, (s-x) *β* → −∞.

**Figure S4:**
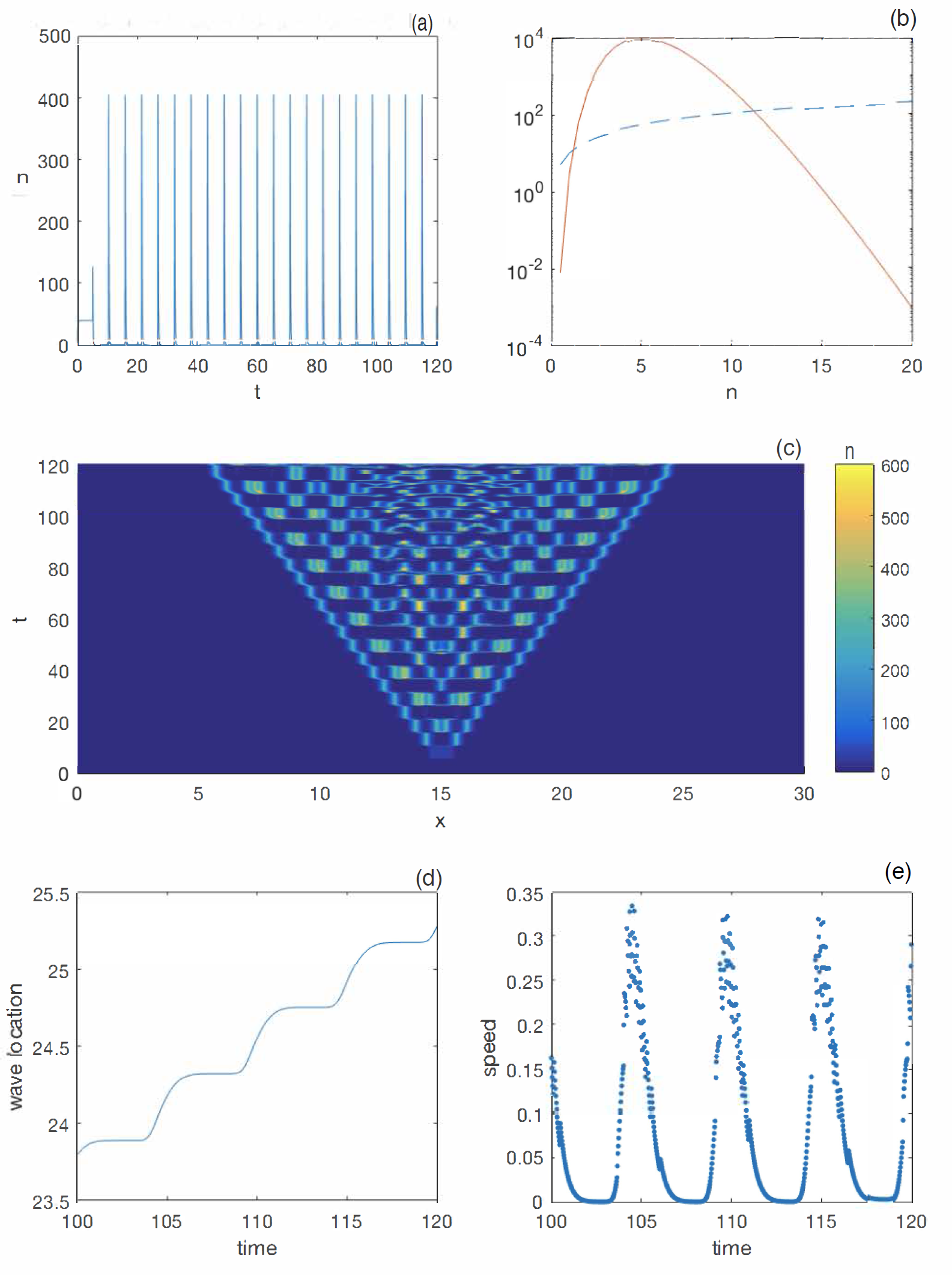
Simulation of model (Eq. 2 from SI Appendix), with *a*_1_ = 20, *a*_2_ = 2, *a*_3_ = 10, *µ* = 10, *τ* = 5, and *D* = 0.1. For these parameters the *undelayed* ordinary differential equation (Eq. 2 from SI Appendix with *D* = 0 and *τ* = 0) exhibits a strong Allee effect, evident by comparing the mortality rate (dashed blue line) to the reproduction rate (solid red curve) in panel (b) (note the logarithmic scale). With delays, the model without movement (Eq. 2 from SI Appendix with *D* = 0) exhibits sharp generational cycles (a). The simulation of the partial functional differential equation (Eq. 2 from SI Appendix; initialized with *n* = 2 for |*x* − 15| ≤ 0.5 and 0 ≤ *t* ≤ *τ*, and *n* = 0 otherwise) exhibits complex dynamics behind the invading fronts (c). These oscillations push the wave forward with a variable speed (d,e).

**Figure S5:**
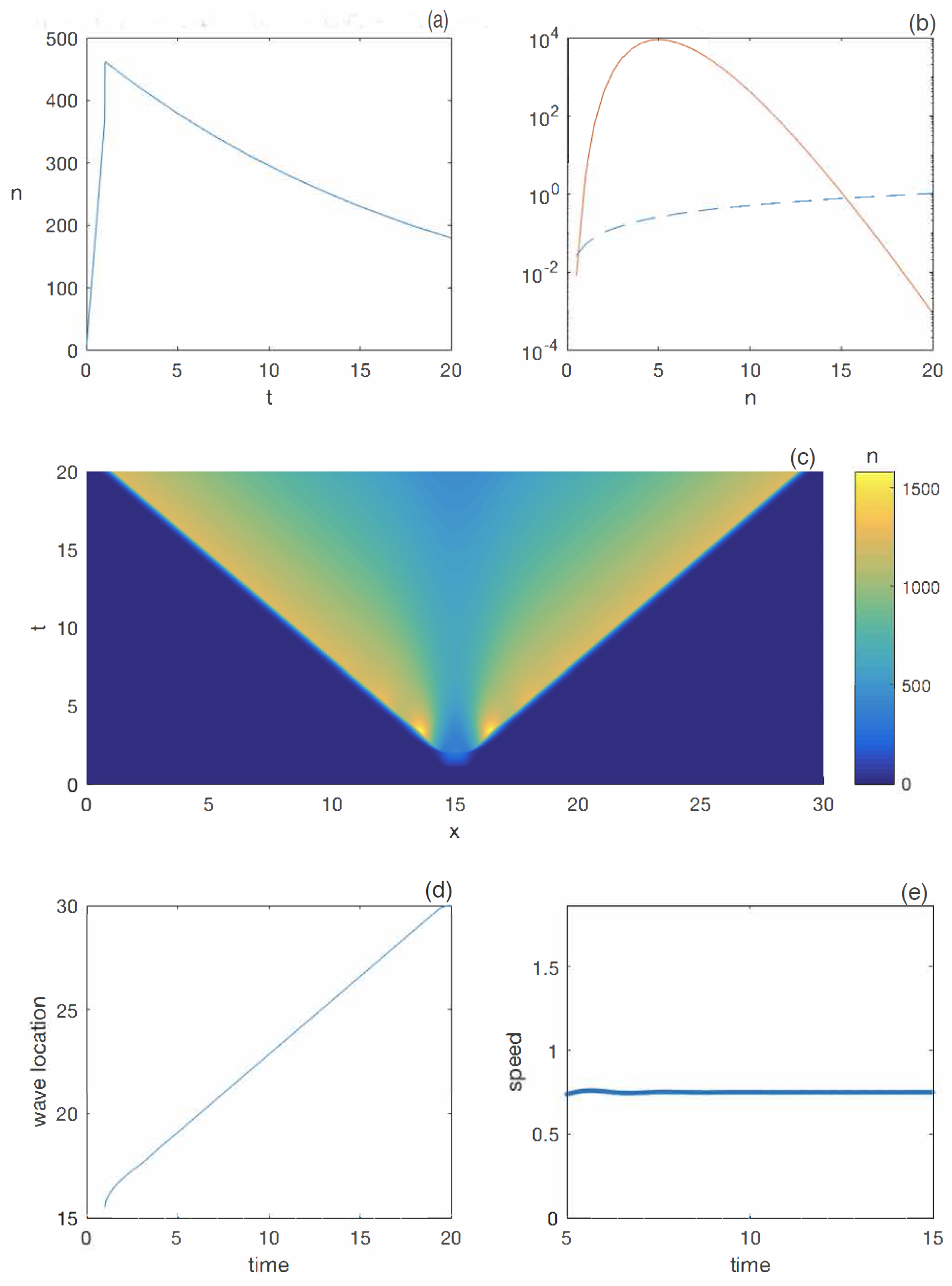
Simulation of model (Eq. 2 from SI Appendix), with *a*_1_ = 20, *a*_2_ = 2, *a*_3_ = 10, *µ* = 0.05, *τ* = 1, and *D* = 0.1. For these parameters the *undelayed* ordinary differential equation (Eq. 2 from SI Appendix with *D* = 0 and *τ* = 0) exhibits a strong Allee effect, evident by comparing the mortality rate (dashed blue line) to the reproduction rate (solid red curve) in panel (b) (note the logarithmic scale). With delays, the model without movement (Eq. 2 from SI Appendix with *D* = 0) exhibits decay to a stable equilibrium (a). The simulation of the partial functional differential equation (Eq. 2 from SI Appendix; initialized with *n* = 2 for |*x* − 15| ≤ 0.5 and 0 ≤ *t* ≤ *τ*, and *n* = 0 otherwise) exhibits simple dynamics behind the invading fronts (c). High population densities behind the wave front push the invasion forward with a constant speed (d,e).

**Figure S6:**
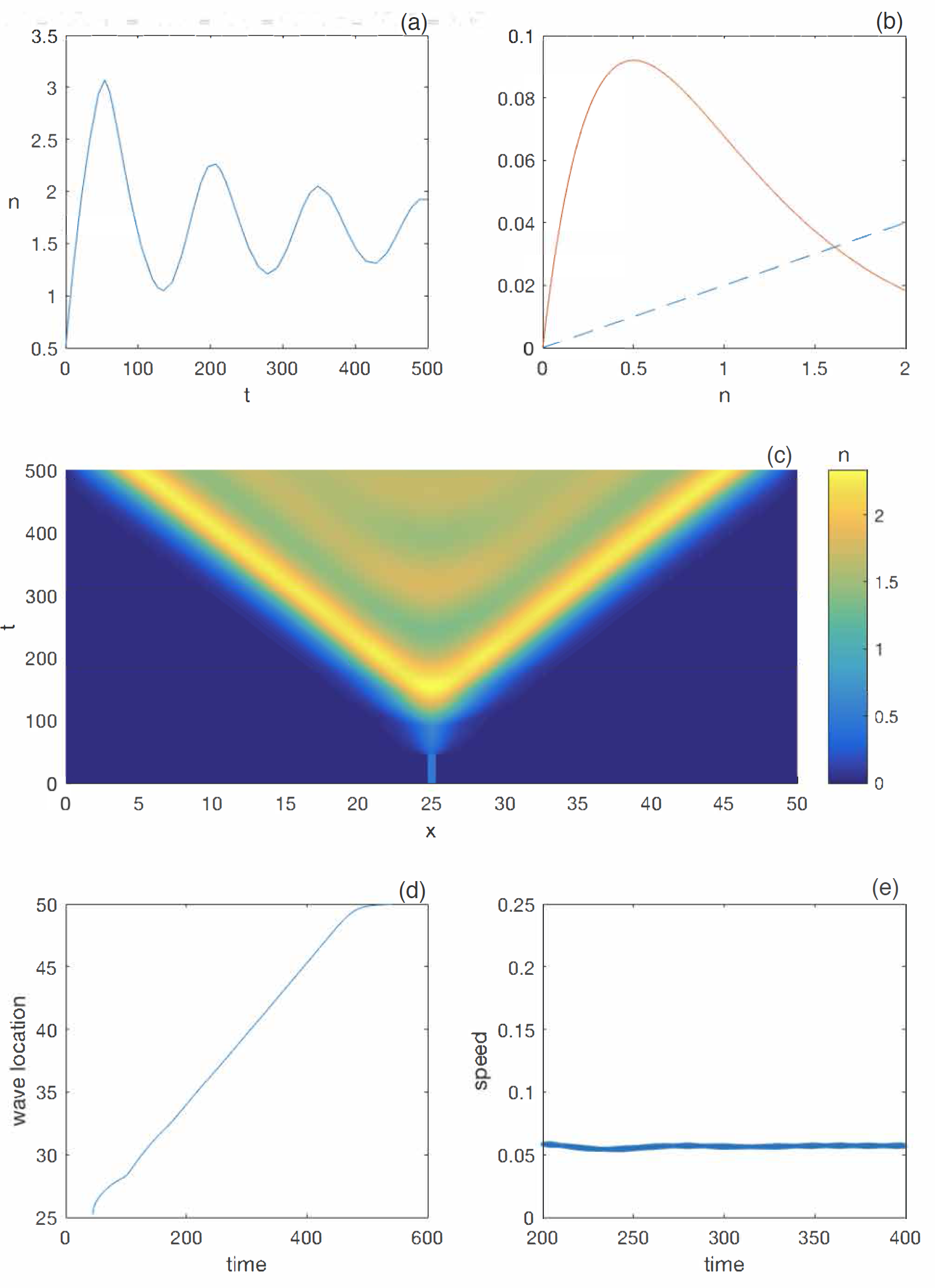
Simulation of model (Eq. 2 from SI Appendix) with *a*_1_ = 0.5, *a*_2_ = 2, *a*_3_ = 1, *µ* = 0.02, *τ* = 45, and *D* = 0.05. For these parameters the *undelayed* ordinary differential equation (Eq. 2 from SI Appendix with *D* = 0 and *τ* = 0) does not exhibit any Allee effect, evident by comparing the mortality rate (dashed blue line) to the reproduction rate (solid red curve) in panel (b) (note the arithmetic scale). With delays, the model without movement (Eq. 2 from SI Appendix with *D* = 0) produces oscillations in population density (a). The simulation of the partial functional differential equation (Eq. 2 from SI Appendix; initialized with *n* = 0.5 for |*x* − 25| ≤ 0.5 and 0 ≤ *t* ≤ *τ*, and *n* = 0 otherwise) exhibits oscillatory dynamics behind the invading fronts (c); however, the wave is “pulled” by growth at low densities, so a constant invasion speed is achieved despite the fluctuations at high densities (d,e).

**Figure S7:**
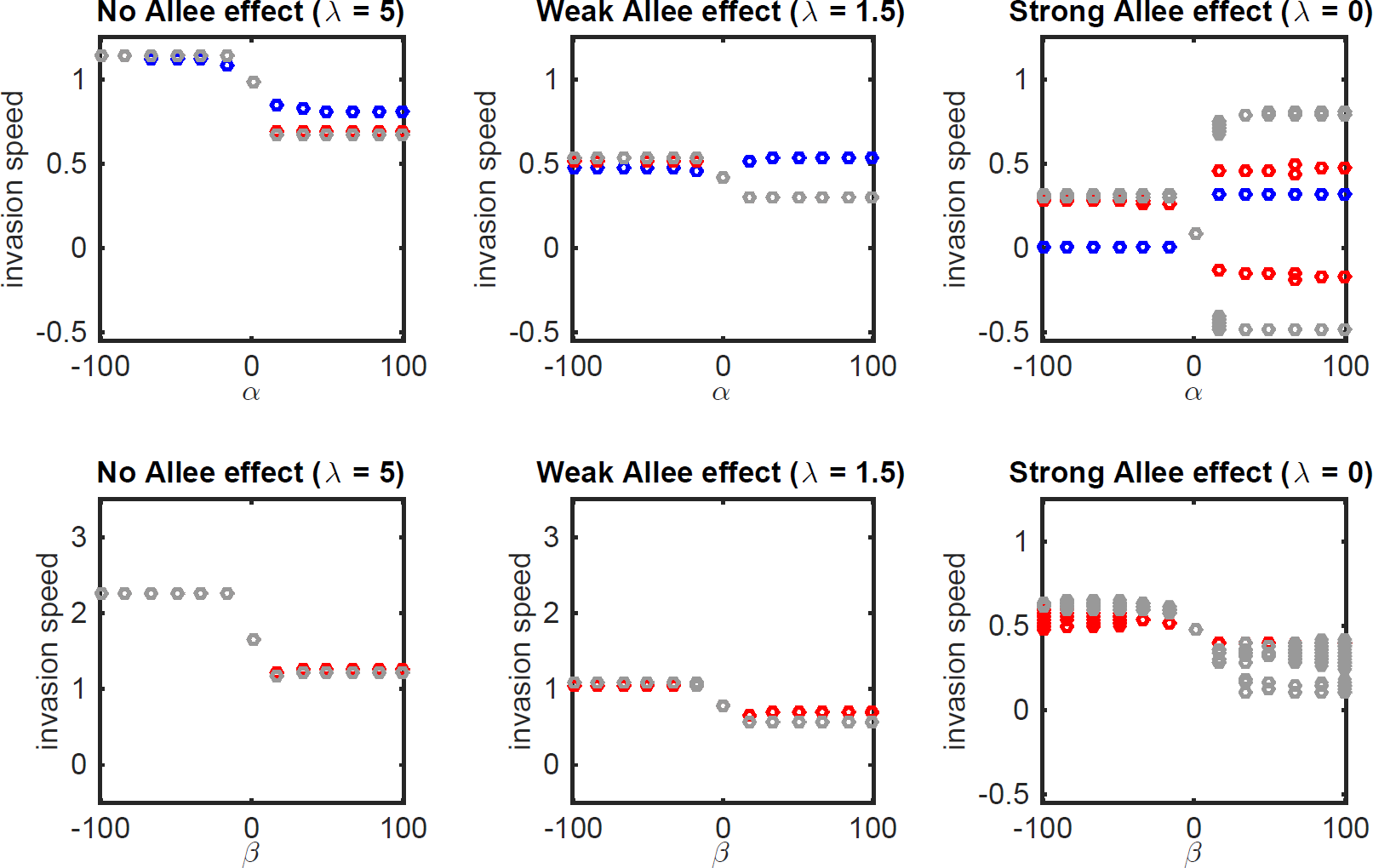
Bifurcation diagram indicating fluctuations in invasion speed across a range of Allee effect strength for the propensity Model – when density dependence alters dispersal propensity (a-c), and the distance model – when density dependence alters dispersal distance (d-f). Here, we also show a range of dispersal thresholds (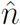) relative to Allee effect threshold used for these models in the text (*a* = 0.2), including 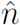 = 0.1 < *a* (blue circles), 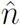 = 0.7 > *a* (red circles), and 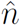 = 0.9 >> *a* (gray circles). All other parameters are the same as Fig. 2. Fluctuations in invasion speed only occur when Allee effects are strong, when the dispersal threshold is high, and when *α* > 0 (propensity model (a-c)) or across a range of positive and negative *β’s* (distance model (d-f)).

**Figure S8:**
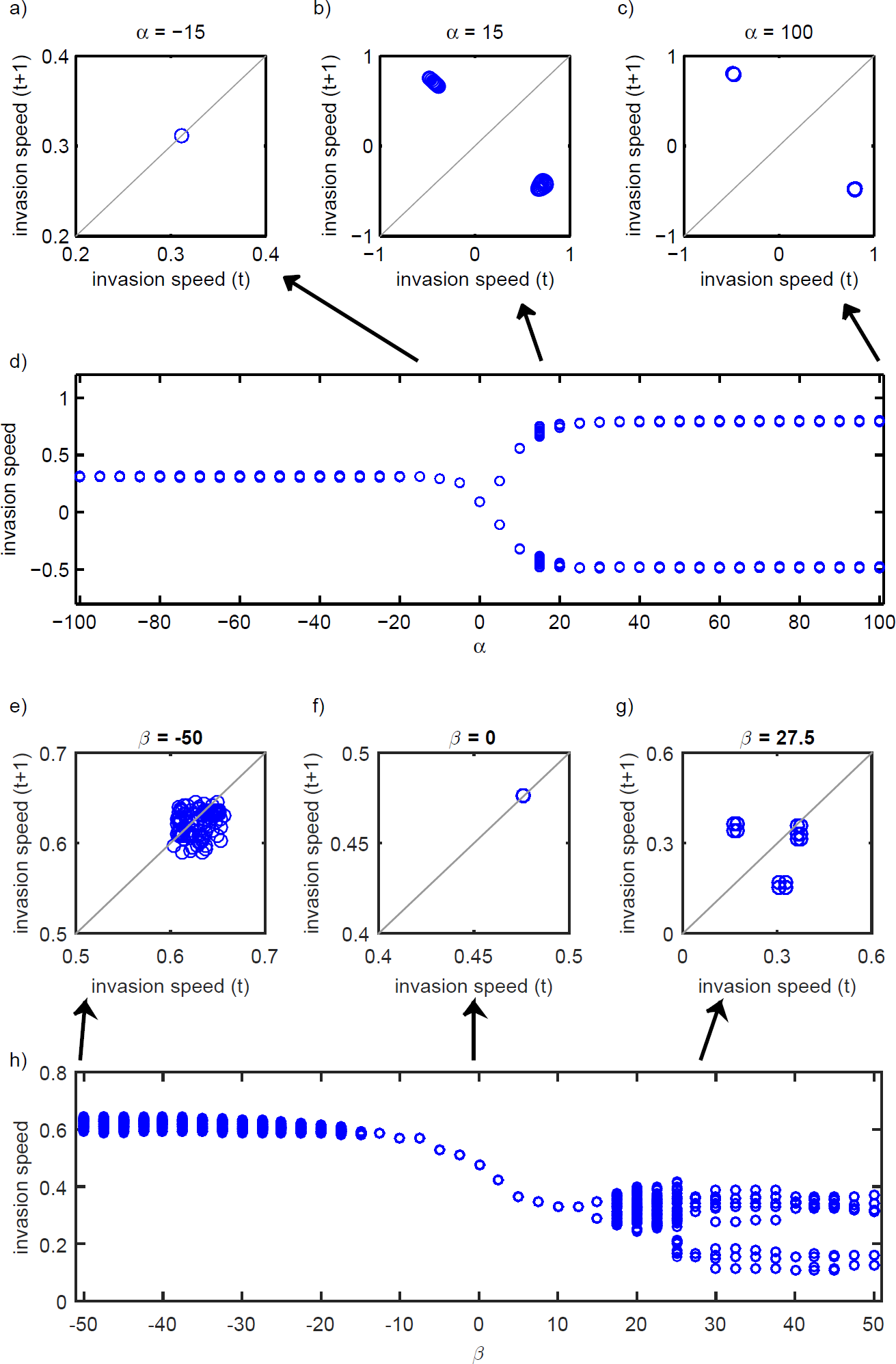
The periodicity of the invasion speed through time for the propensity model (Allee effects and density-dependent dispersal propensity; a-d) and distance model (Allee effects and density-dependent dispersal *distance*; e-h). In panels a-c and e-g, the wave position is plotted at time *t* vs time *t* + 1. In panels d and h, the range of invasion speeds represents the amplitude of fluctuations. For each parameter value, the invasion speed for the previous 100 time steps are plotted. When points appear as hollow points, the same invasion speed is being plotted over itself many times. For the propensity Model, when fluctuating, the wave speed is nearly always periodic across values of the Allee effect threshold *a*. At small values of *α* the invasion speed is constant (a), at small positive *α* the invasion speed fluctuates in a quasi-periodic fashion (b), and most positive *α* values, for example *α* = 100 (c), the wave speed is periodic. Here, 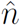 = 0.9, *λ* = 0, *σ*^2^ = 0.25, *p*_0_ = 0.05, *p*_*max*_ = 1, and *a* = 0.2. For the distance model, we demonstrate that the invasion speed appears to be more chaotic for some negative values of the density-dependent dispersal threshold (*β*) (e), is constant for some values of *β*(f), and has a quasi-periodic attractor for some positive values of *β*(g). Here, 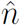 = 0.9, *λ* = 0, 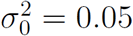, 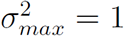, and *α* = 0.2.

## SI Appendix

Here we construct a continuous-time model that we conjecture produces variable-speed invasions. We begin with a modification of a delay-differential equation model used by Gurney et al. (1) to study the dynamics of “Nicholson’s blowflies:” 
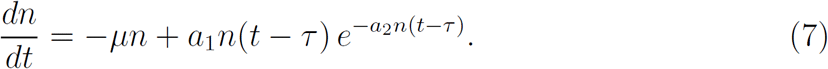

In this model, *n* is the population size of mature animals, and *τ* is the maturation time. The change in the adult population size is due to constant per captia mortality (at rate *µ*) and recruitment of juveniles, born *τ* time units ago, into the adult class. The per captia birth rate at low density (*a*_1_) is reduced (exponentially at the rate *a*_2_) at larger population densities. This model produces large swings in adult population size when the maturation time is sufficiently large (1).

We modify the model (7) to include the potential for a strong Allee effect (when the parameter *a*_3_ > 1) and to include the random movement of adults via diffusion:

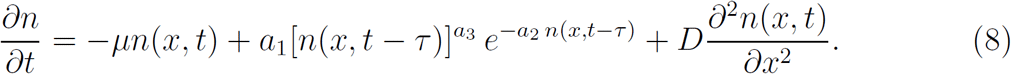

Immature individuals are assumed to be sedentary.

The special case of model (8) with *a*_3_ = 1 (without Allee effects) has been thoroughly studied (see, e.g., Lin et al. (2) and Solar and Trofimchuk (3) and references therein). The dynamics of this model in this case can be quite complex behind the leading invasion front, but, for biologically realistic initial conditions solutions, solutions exhibit an asymptotically constant spreading speed.

Much less is know about the dynamics of equation (8) when *a*_3_ > 1, but the model would seem to have the features necessary to generate variable invasion speed. Density-dependent reproduction, along with the maturation time delay, induce population fluctuations at high density, and the Allee effect should generate a pushed wave. Our numerical simulations suggest that this is indeed the case (Fig. S4). When, in contrast, the population dynamics converge to an equilibrium point behind the invasion front, the invasion speed is eventually constant, even in the presence of an Allee effect (Fig. S5). In the absence of Allee effects (*a*_3_ = 1), simulations of the model (8) produce constant speed invasions (Fig. S6), even if there are oscillatory dynamics behind the front, in agreement with prior theory.

